# Development of a CRISPR-Cas12a Assay for Detection of the P*nimB*^G^ Mutation Associated with Heme-Dependent Metronidazole Resistance in *Clostridioides difficile*

**DOI:** 10.64898/2026.10.06.756461

**Authors:** Nicola Szadna, Cayden Massy, Abiola O. Olaitan

**Author notes:** Corresponding author: Abiola O. Olaitan, Department of Biology, University of Waterloo, Waterloo, ON, Canada.

## Abstract

**Objective:** Heme-dependent metronidazole (MTZ) resistance has emerged in *Clostridioides difficile* and is primarily mediated by a T-to-G single-nucleotide polymorphism (SNP) in the promoter of *nimB* (P*nimB*^G^). We aimed to develop a CRISPR-Cas12a-based assay for rapid detection of the P*nimB*^G^ mutation associated with MTZ resistance.

**Methods:** A series of primers and CRISPR RNAs (crRNAs) were designed and optimized to identify the most effective combination for discriminating between P*nimB*^G^ (MTZ-resistant) and P*nimB*^T^ (MTZ-susceptible) alleles. Detection was evaluated by visual fluorescence using blue-light and UV imaging systems, as well as by quantitative fluorescence measurements.

**Results:** The CRISPR-Cas12a assay accurately differentiated P*nimB*^G^ and P*nimB*^T^ alleles in *C. difficile*. In a collection of clinical isolates, the assay correctly identified all 11 MTZ-susceptible isolates carrying the P*nimB*^T^ allele and all 10 MTZ-resistant isolates carrying the P*nimB*^G^ allele. Fluorescence signals were readily detected both visually and quantitatively. Furthermore, colony PCR using crude colony lysates yielded results comparable to those obtained with purified genomic DNA, enabling rapid colony-based detection without conventional DNA extraction.

**Conclusions:** The CRISPR-Cas12a assay enables rapid molecular detection of the P*nimB*^G^ mutation associated with heme-dependent MTZ resistance in *C. difficile*. This approach could complement conventional susceptibility testing for MTZ resistance detection.

## Introduction

*Clostridioides difficile* infection (CDI) is a leading cause of healthcare-associated diarrhea worldwide and is associated with substantial morbidity and healthcare costs [1]. Metronidazole (MTZ), vancomycin (VAN), and fidaxomicin (FDX) are the primary antibiotics used to treat CDI. Although MTZ use has declined because of reduced clinical effectiveness and increasing resistance [2]

Heme-dependent MTZ resistance has recently emerged and is a common MTZ resistance mechanism in clinical *C. difficile* isolates [3,4]. Resistant strains exhibit a ≥4-fold increase in MTZ minimum inhibitory concentration (MIC) in the presence of heme compared to the absence of heme, and this phenotype has been associated with poor treatment outcomes [5]. Heme-dependent MTZ resistance is primarily mediated by a T-to-G substitution located 43 nt upstream of the *nimB* start codon in the promoter region (P*nimB*^G^), which drives constitutive *nimB* expression [6]. A recent population-genomics study identified the P*nimB*^G^ mutation in 19.5% of publicly available *C. difficile* isolates across diverse ribotypes and regions, demonstrating the widespread dissemination of this resistance mechanism [7].

Detection of heme-dependent MTZ resistance currently relies on phenotypic susceptibility testing or identification of the P*nimB*^G^ mutation by sequencing. However, phenotypic testing requires careful preparation and handling of heme-containing media, as heme degradation caused by light exposure can lead to false-negative susceptibility results [3,8], whereas sequencing-based approaches may not be readily accessible in all settings.

CRISPR-Cas12a has emerged as a powerful molecular diagnostic tool for the detection of antimicrobial resistance determinants and bacterial pathogens [9–11]. In this system, a guide CRISPR RNA (crRNA) directs Cas12a to a complementary DNA target, triggering collateral cleavage of fluorophore-quencher-labeled single-stranded DNA reporters and generating a detectable fluorescence signal [12]. Importantly, the ability of Cas12a enzyme to discriminate single-nucleotide polymorphisms (SNP) makes it particularly well suited for detection of the P*nimB*^G^ mutation.

Here, we developed and optimized a CRISPR-Cas12a assay for rapid detection of the P*nimB*^G^ mutation and evaluated its ability to distinguish MTZ-resistant from MTZ-susceptible *C. difficile* isolates.

## Materials and Methods

### Bacterial strains

*Clostridioides difficile* strains R20291 and CD196 were used as representative MTZ-resistant and MTZ-susceptible strains, respectively. Additional clinical isolates comprising both MTZ-resistant and MTZ-susceptible strains were obtained from multiple sources, including the Texas Medical Center Biodefense and Emerging Infections (BEI) Resources, USA and Italy. Bacteria were cultured in pre-reduced brain heart infusion (BHI) broth or on BHI agar and incubated anaerobically at 37°C in a Whitley A35 anaerobic workstation containing 10% H_2_, 5% CO_2_, and 85% N_2_ (Don Whitley Scientific). MTZ MICs were determined as previously described [6], except for strain JH7, for which the previously reported MIC value was used. P*nimB* promoter SNPs which have been previously characterized and reported were obtained from published data [6]. P*nimB* SNPs for additional isolates that were not previously characterized were sequenced using the Sanger method.

### PCR primer and crRNA design for SNP detection

Because the P*nimB*^G^ SNP occurs at a fixed genomic position, Cas12a-based detection requires the presence of a suitable protospacer adjacent motif (PAM; TTTN) near the SNP. To identify the optimal PAM position for SNP discrimination, PAM sites were introduced at varying distances relative to the SNP through PCR primer design (Table S1). Six forward primers and a common reverse primer were designed to generate amplicons in which the PAM site was positioned 0-5 nucleotides (nt) from the SNP on the coding strand. In principle, the non-coding strand could also be targeted, thereby expanding the number of candidate PAM positions [13]. Similarly, six crRNAs were designed to match each PAM position. Similar to previous study [13], each crRNA consisted of a universal 19-nucleotide direct-repeat sequence (5′-AAUUUCUACUGUUGUAGAU-3′) and a 17-nucleotide spacer sequence complementary to the target region immediately downstream of the PAM site (Table S1). All primers were synthesized by Sigma-Aldrich, and crRNAs were synthesized by Thermo Fisher Scientific. A fluorescent reporter consisting of a random 12-nt single-stranded DNA oligonucleotide labeled with ROX at the 5′ end and BHQ1 at the 3′ end (ROX-5′-GAGCCTGACATG-3′-BHQ1) was synthesized by Integrated DNA Technologies (IDT, USA).

### crRNA screening and optimization

Genomic DNA (gDNA) was extracted from overnight cultures using the EZ-10 Spin Column Bacterial Genomic DNA Miniprep Kit (Bio Basic, Canada; Cat. No. BS423) or the DNeasy UltraClean Microbial Kit (Qiagen) according to the manufacturers’ instructions. For direct colony DNA extraction, colonies grown anaerobically on BHI agar at 37°C overnight were suspended in 50 μL TES buffer (50 mM Tris-HCl [pH 8.0], 5 mM EDTA, and 50 mM NaCl). Cell suspensions were heat-lysed at 95°C for 10 min and centrifuged at 10,000 rpm for 1 min. The resulting supernatants were used directly as DNA templates for colony PCR.

PCR amplification was performed in 25-μL reaction volumes using GB-AMP PaCeR HP High-Fidelity Master Mix (GeneBioSystems, Canada) with 2 μL of DNA template per reaction. Primer sequences and PCR thermocycling conditions are provided in Tables S1 and S2, respectively. Following amplification, 2 μL of unpurified PCR product was added directly to the CRISPR-Cas12a reaction mixture (Table S3). EnGen Lba Cas12a (Cpf1) was purchased from New England Biolabs (NEB, Canada). CRISPR-Cas12a reactions were incubated at 37°C for 15 min, followed by enzyme inactivation at 65°C for 10 min. Fluorescence signals were visualized using an Edvotek TruBlu 2 Blue/White Transilluminator and a Bio-Rad ChemiDoc MP Imaging System. Quantitative fluorescence measurements were performed using Nunc 96-well black microplates (Cat. No. 165305; Thermo Fisher Scientific) and a Synergy H1 microplate reader (Agilent BioTek) with excitation and emission wavelengths of 575 nm and 602 nm, respectively. Fluorescence visualization and measurements were conducted within 1 h of reaction completion to minimize signal loss. Negative controls consisted of PCR reactions containing nuclease-free water or TES buffer in place of the DNA template. Fluorescence values were normalized by subtracting the corresponding negative-control signal from each sample. Samples with fluorescence values equal to or below the negative control were assigned a value of 1 relative fluorescence unit (RFU).

## Results

### Optimization of PCR primers and crRNAs for the detection of P*nimB*^G^

Using the MTZ-susceptible strain CD196 and the MTZ-resistant strain R20291, six PCR amplicon/crRNA combinations targeting SNP positions 0-5 bp from the PAM-proximal end of the protospacer were evaluated. CrRNAs with the SNP positioned 1-3 bp from the PAM-proximal end generated the strongest fluorescence signals, as determined by both blue-light/UV visualization and quantitative fluorescence measurements (Fig. 1a, b). Notably, crRNA-4, in which the SNP was positioned 3 bp from the PAM-proximal end, produced the highest signal-to-noise ratio, yielding a 4,034-fold difference in fluorescence between P*nimB*^G^ and P*nimB*^T^ templates (Fig. 1c). Therefore, the *nimB*-4 forward/reverse primer pair and crRNA_*nimB*-4 were selected for all subsequent analyses.

**Fig. 1.**
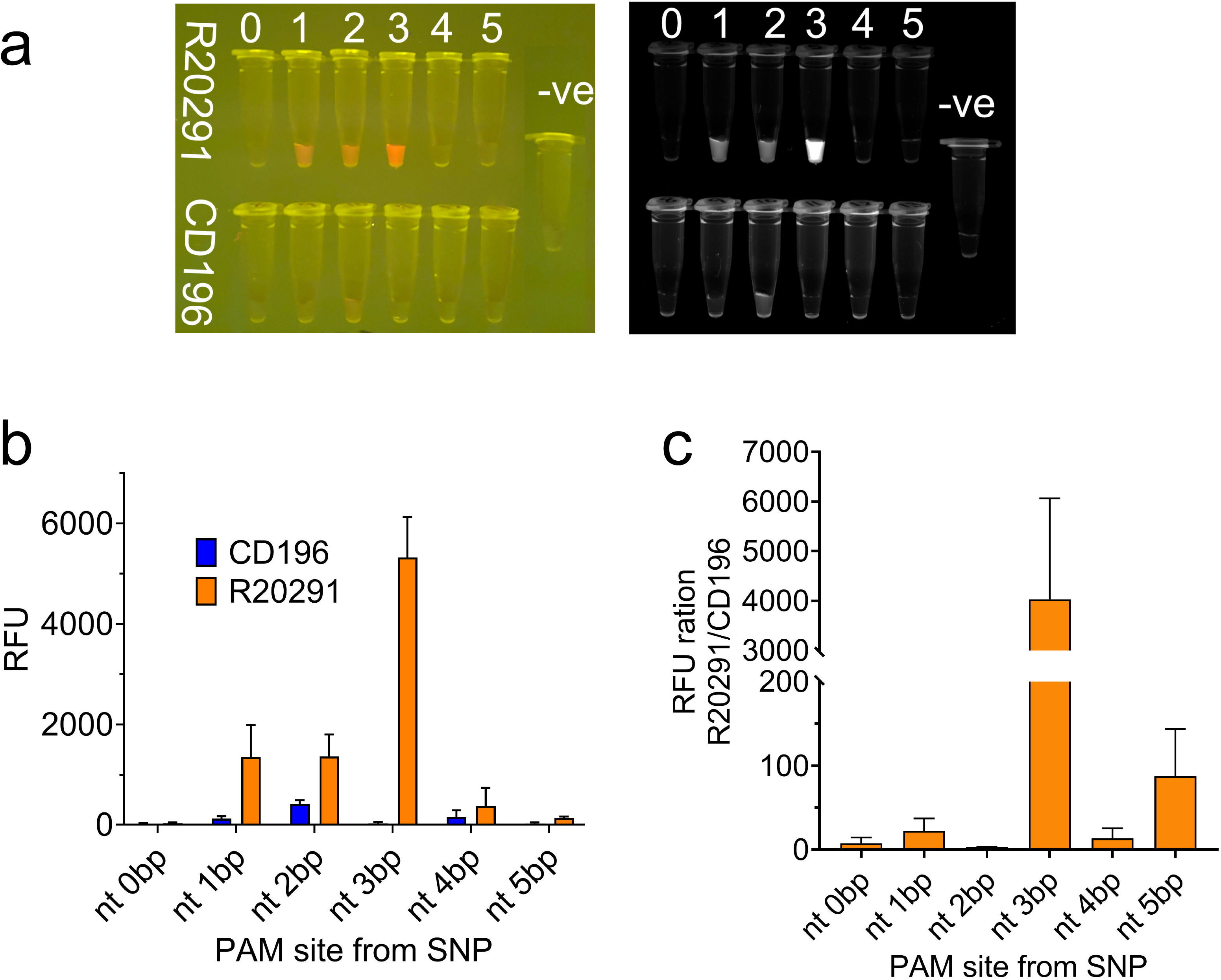
Optimization of P*nimB*^G^ detection. Genomic DNA from CD196, an MTZ-susceptible strain carrying the wild-type *PnimB* promoter (P*nimB*^T^), and R20291, an MTZ-resistant strain carrying P*nimB*^G^, was used for CRISPR-Cas12a assay optimization. **(a)** Fluorescence visualization under blue-light and UV illumination using a DNA gel imaging system. **(b)** Relative fluorescence units (RFU) obtained for CD196 and R20291 using crRNAs with the target SNP positioned 1, 2, or 3 nt downstream of the PAM. **(c)** RFU ratio between R20291 and CD196 for each crRNA design. For assay optimization, three independent technical replicates of the CRISPR-Cas12a reaction were performed using the same PCR product. Data are presented as mean ± SEM.

### Detection of P*nimB*^G^ in Clinical *C. difficile* Isolates Using the CRISPR-Cas12a Assay

A collection of 23 *C. difficile* isolates with previously characterized MTZ susceptibility profiles, comprising 11 MTZ-susceptible and 12 MTZ-resistant isolates, was screened using the optimized CRISPR-Cas12a assay. All 11 MTZ-susceptible isolates possessed the wild-type P*nimB*^T^ allele, whereas 10 of the 12 MTZ-resistant isolates carried the P*nimB*^G^ allele (Fig. 2a). The remaining two MTZ-resistant isolates lacked the P*nimB*^G^ allele.

**Fig. 2.**
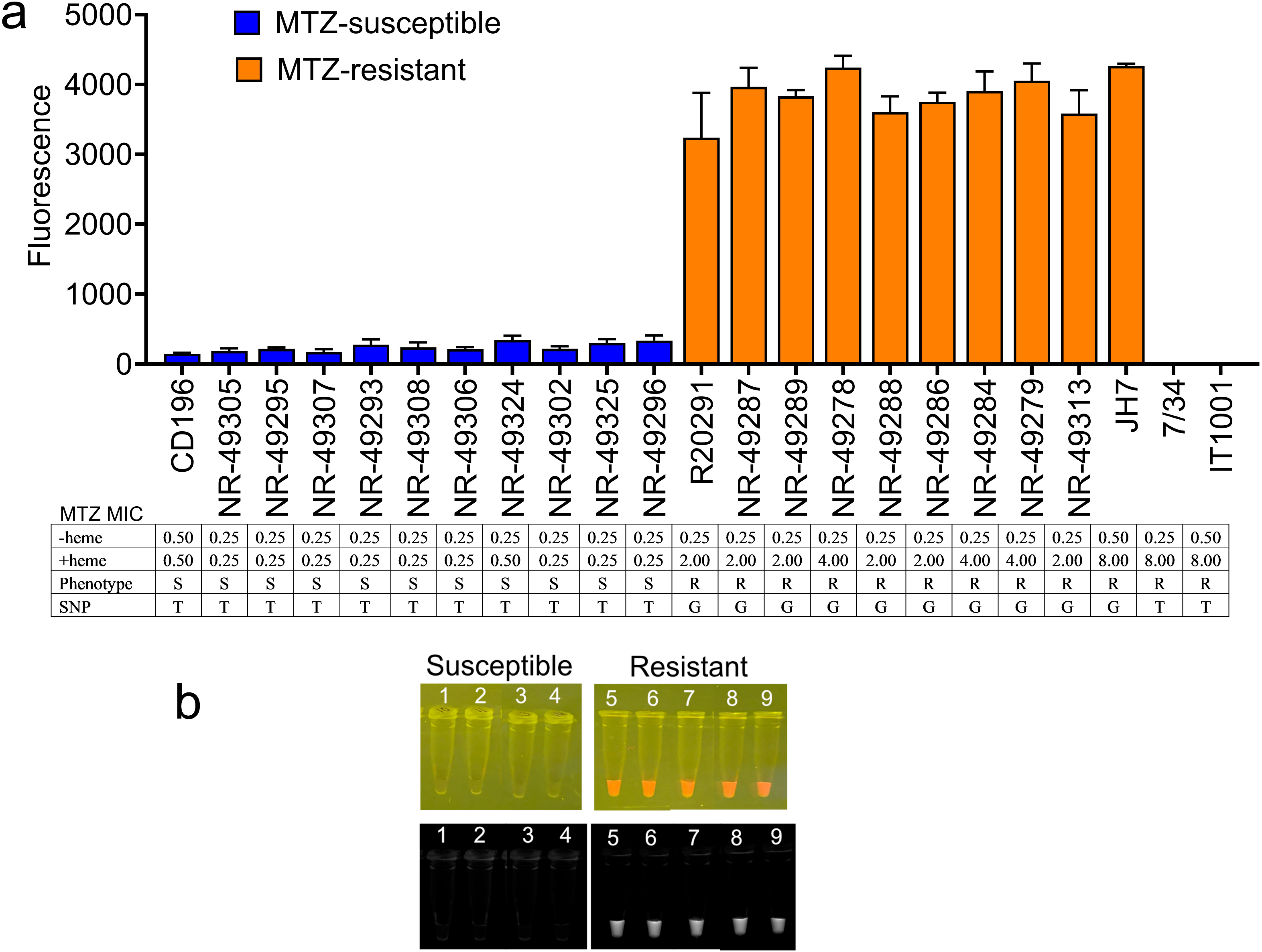
Detection of P*nimB*^G^ in clinical isolates. **(a)** CRISPR-Cas12a detection of P*nimB*^G^ in clinical *C. difficile* isolates. MTZ-susceptible isolates carrying the wild-type promoter (P*nimB*^T^) showed no detectable fluorescence, whereas MTZ-resistant isolates carrying P*nimB*^G^ generated strong fluorescence signals. Isolate genotypes and phenotypes are provided in the accompanying table. **(b)** Representative blue-light images from an independent experiment performed under the same conditions. 1. NR-49307, 2. NR-49293, 3. NR-49308, 4. NR-49306, 5. NR-49287, 6. NR-49289, 7. NR-49278, 8. NR-49288, 9. NR-49286. Data are presented as mean ± SEM from three biological replicates.

Consistent with their P*nimB* genotypes, none of the 11 MTZ-susceptible isolates produced fluorescence detectable by visual inspection, with relative fluorescence unit (RFU) values ranging from 147 to 346 (Fig. 2a, b). In contrast, all 10 MTZ-resistant isolates carrying the P*nimB*^G^ allele produced strong visible fluorescence signals, with RFU values ranging from 3,237 to 4,265 (Fig. 2a, b). No detectable fluorescence was observed for the two MTZ-resistant isolates lacking the P*nimB*^G^ allele.

Overall, P*nimB* genotypes determined by the CRISPR-Cas12a assay were fully concordant with whole-genome or Sanger sequencing results (Fig. 2a).

### P*nimB*^G^ Detection Using Colony PCR

To evaluate whether crude colony-derived DNA could be used in place of purified genomic DNA (gDNA), four MTZ-susceptible and four MTZ-resistant *C. difficile* isolates were tested using the CRISPR-Cas12a assay. Crude DNA extracts prepared directly from bacterial colonies were used as templates for PCR amplification, followed by CRISPR-Cas12a detection.

None of the four MTZ-susceptible isolates produced detectable fluorescence (Fig. 3). In contrast, all four MTZ-resistant isolates generated strong fluorescence signals, determined by both visual detection under blue-light illumination and quantitative fluorescence measurements (Fig. 3). Overall, colony PCR yielded P*nimB*^G^ detection profiles similar to those obtained using purified gDNA templates.

**Fig. 3.**
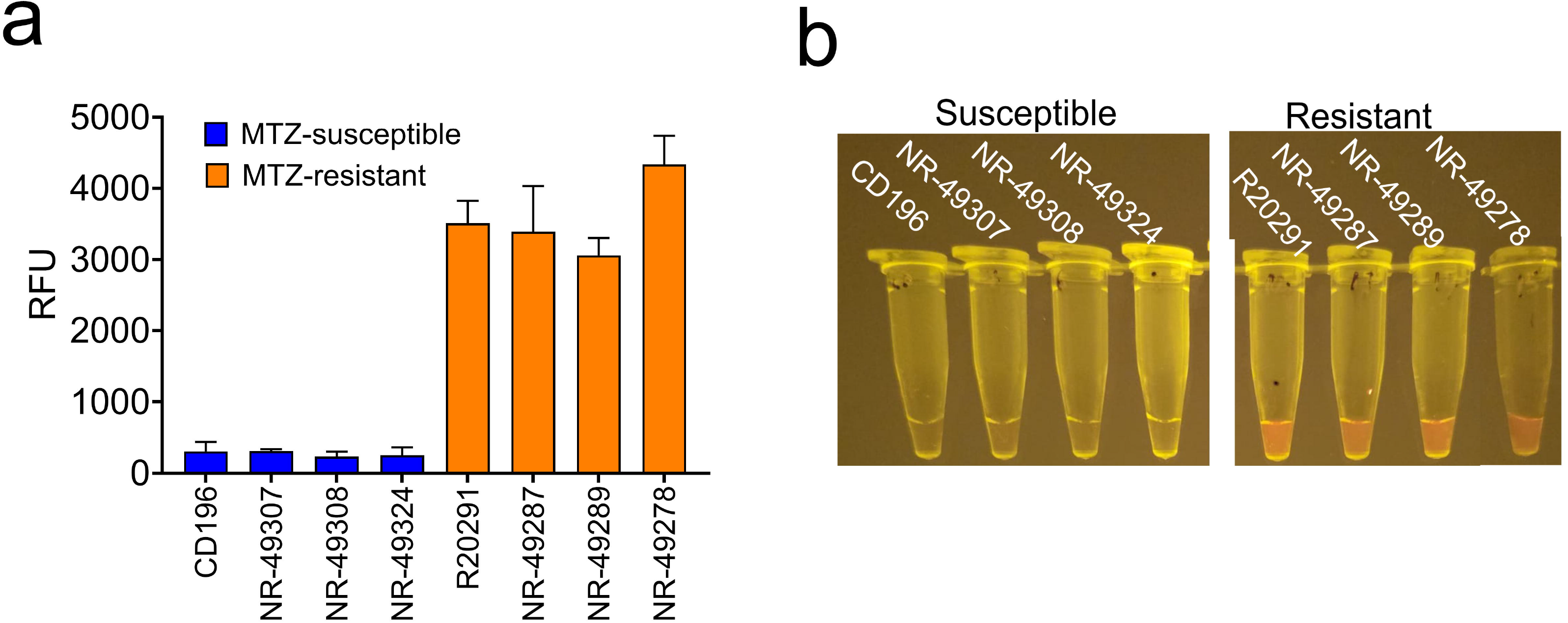
Detection of P*nimB*^G^ using crude colony-derived DNA extracts. **(a)** Crude DNA extracts prepared directly from colonies of selected clinical *C. difficile* isolates were used as templates for PCR amplification followed by CRISPR-Cas12a detection. MTZ-resistant isolates carrying P*nimB*^G^ generated strong fluorescence signals, whereas MTZ-susceptible isolates carrying the wild-type promoter (P*nimB*^T^) showed no detectable fluorescence. **(b)** Representative blue-light fluorescence images of the isolates shown in panel (a). Data are presented as mean ± SEM from three biological replicates.

## Discussion

Heme-dependent MTZ resistance mediated by the P*nimB*^G^ promoter mutation has now been reported in clinical *C. difficile* isolates from North America, Europe, and Asia [6,8,14,15]. Genome-wide analyses have further demonstrated that this mutation is distributed across all continents, including Africa [7]. The identification of P*nimB*^G^ as the primary determinant of this resistance phenotype provides an opportunity for molecular diagnosis, particularly given the challenges associated with phenotypic susceptibility testing [3,4,8,15].

In this study, we developed a CRISPR-Cas12a assay for the detection of P*nimB*^G^, the mutation associated with heme-dependent MTZ resistance. A critical step in assay development was identifying the optimal SNP position relative to the PAM site for Cas12a discrimination. We found that positioning the SNP three nucleotides from the PAM-proximal end of the protospacer provided the greatest discrimination between P*nimB*^G^ and P*nimB*^T^. Using this optimized design, all P*nimB*^G^ -positive MTZ-resistant isolates were correctly identified, whereas no signal was detected in susceptible isolates carrying the wild-type P*nimB*^T^ allele. The assay did not detect the two MTZ-resistant isolates lacking P*nimB*^G^, consistent with previous observations that a minority (∼10%) of heme-dependent resistant strains possess alternative, currently unknown resistance mechanisms [6]. Because the majority of MTZ-resistant isolates carry P*nimB*^G^, the assay is expected to identify most cases of heme-dependent MTZ resistance.

An additional advantage of the assay is its simplicity and flexibility. Cas12a-generated fluorescence could be visualized using standard UV/blue-light imaging systems in addition to a fluorescence plate reader, enabling implementation in laboratories without specialized fluorescence detection equipment. Furthermore, colony PCR yielded results comparable to those obtained using purified gDNA, eliminating conventional DNA extraction and reducing assay turnaround time.

Resistance to other anti-*C. difficile* agents, including vancomycin and fidaxomicin, has also been associated with resistance-conferring SNPs in *vanR*/*vanS* and *rpoB*, respectively [7,16,17], suggesting that CRISPR-based assays may have broader applications for the detection of antimicrobial resistance in *C. difficile*. However, our preliminary attempts to target the *vanR* 343A>G mutation, which results in the VanR Thr115Ala substitution and vancomycin resistance in *C. difficile*, were unsuccessful. This observation highlights the current limitation of Cas12a, as it may exhibit reduced discrimination for certain SNP targets. The use of alternative Cas12a variants or other CRISPR-associated nucleases with greater single-nucleotide discrimination may provide a means to overcome this limitation.

Furthermore, unlike P*nimB*^G^-mediated MTZ resistance, resistance to vancomycin and fidaxomicin may be associated with a range of alternative resistance-conferring mutations in the same gene or in different genes, making it challenging to develop a single assay capable of detecting all relevant mutations. One strategy to address this challenge would be to develop a panel of complementary assays targeting these additional SNPs.

Although MTZ is no longer a first-line therapy for CDI in North America and Europe, it remains widely used in pediatric patients, for non-severe CDI, and in many low- and middle-income countries [18,19]. Given that the P*nimB*^G^ mutation has been detected in strains from all continents, rapid identification of MTZ resistance remains clinically relevant. In the face of increasing antimicrobial resistance and a diminishing antibiotic development pipeline, susceptibility-guided therapy, which is rarely performed in routine CDI management, may represent an important yet underutilized antimicrobial stewardship approach. Rapid molecular assays capable of detecting MTZ resistance could support treatment decisions by identifying patients unlikely to benefit from MTZ therapy. Combined with our recent finding that artemisinin restores MTZ susceptibility in resistant isolates, these findings underscore the potential of improved diagnostics and adjunctive therapies to preserve the effectiveness of existing anti-*C. difficile* agents.

In conclusion, we developed a CRISPR-Cas12a assay that enables rapid detection of P*nimB*^G^-mediated MTZ resistance in *C. difficile* and could serve as a practical molecular diagnostic tool alongside the conventional susceptibility testing.

## Acknowledgements

The authors thank Dr. Julian Hurdle (Texas A&M Health Science Center, Houston, TX, USA), Dr. Kevin Garey (University of Houston, Houston, TX, USA), and Dr. Patrizia Spigaglia (Istituto Superiore di Sanità, Rome, Italy) for providing clinical *C. difficile* isolates. The authors also acknowledge BEI Resources, National Institute of Allergy and Infectious Diseases, National Institutes of Health, for supplying additional isolates used in this study.

This work was supported by the University of Waterloo through funding awarded to A.O.O. The funder had no role in study design, data collection and interpretation, or the decision to submit the work for publication.

