## Supplementary material for "Development of a CRISPR-Cas12a Assay for Detection of the P*nimB*^G^ Mutation Associated with Heme-Dependent Metronidazole Resistance in *Clostridioides difficile*": Table S1, S2, S3

**Table S1**. Primers and crRNAs used for optimization of P*nimB*^G^ detection by the CRISPR-Cas12a assay.

| Target sequence 5’-3’ | PAM Sequence | Primers 5’-3’ | | crRNA 5’-3’ |
| --- | --- | --- | --- | --- |
| ATGT**G**CTAAAATTTAATTTAA | TTTT | nim-A1F | AACATAGCATAATCTTTATAA**TTTT** | AAUUUCUACUGUUGUAGAU  GCUAAAAUUUAAUUUAA |
|  |  | nim-R | AATACTTCAACAGTATCTTCTTTTGT |  |
| AATGT**G**CTAAAATTTAATTTA | TTTG | nim-A2F | AACATAGCATAATCTTTATA**TTTG**T | AAUUUCUACUGUUGUAGAU  UGCUAAAAUUUAAUUUA |
|  |  | nim-R | Ditto |  |
| AAATGT**G**CTAAAATTTAATTT | TTTT | nim-A3F | AACATAGCATAATCTTTAT**TTTT**GT | AAUUUCUACUGUUGUAGAU  GUGCUAAAAUUUAAUUU |
|  |  | nim-R | Ditto |  |
| TAAATGT**G**CTAAAATTTAATT | TTTA | nim-A4F | AACATAGCATAATCTTTA**TTTA**TGT | AAUUUCUACUGUUGUAGAU  UGUGCUAAAAUUUAAUU |
|  |  | nim-R | Ditto |  |
| ATAAATGT**G**CTAAAATTTAAT | TTTA | nim-A5F | AACATAGCATAATCTTT**TTTA**ATGT | AAUUUCUACUGUUGUAGAU  AUGUGCUAAAAUUUAAU |
|  |  | nim-R | Ditto |  |
| TATAAATGT**G**CTAAAATTTAA | TTTA | nim-A6F | AACATAGCATAATCTT**TTTA**AATGT | AAUUUCUACUGUUGUAGAU  AAUGUGCUAAAAUUUAA |
|  |  | nim-R | Ditto |  |

Bold text indicates the SNP position in the target sequence and the PAM sequence in the primer. Underlined text indicates the target sequence and the 17-nt crRNA spacer sequence complementary to the target region.

**Table S2**. PCR thermocycling conditions used for amplification of the P*nimB* promoter region prior to CRISPR-Cas12a detection.

| Temperature | Time |
| --- | --- |
| Initial: 95°C | 3 mins |
| Denaturation: 95°C | 15 secs |
| Annealing: 48°C | 15 secs |
| Extension: 72°C | 15 secs |
| Final Extension: 72°C | 5 mins |
| Hold: 4°C | ꝏ |

**Table S3**. Composition of CRISPR-Cas12a reaction mixtures.

| Reagent | Volume (µL) per reaction |
| --- | --- |
| Nuclease free water | 10 |
| NEbuffer 2.1 | 2 |
| LbCas12a nuclease (0.15 mg/mL) | 1 |
| Probe (10 µM) | 1 |
| crRNA (2.5 µM) | 4 |
| PCR product | 2 |
| Total = | 20 µL |
